# A genome-wide association study identifies that the GDF5 and COL27A1 genes are associated with knee pain in UK Biobank (N = 171, 516)

**DOI:** 10.1101/525147

**Authors:** Weihua Meng, Mark J Adams, Colin NA Palmer, The 23andMe Research Team, Jingchunzi Shi, Adam Auton, Kathleen A. Ryan, Joanne M. Jordan, Braxton D. Mitchell, Rebecca D. Jackson, Michelle S. Yau, Andrew M McIntosh, Blair H Smith

## Abstract

**Objective:** Knee pain is one of the most common musculoskeletal complaints that brings people to medical attention. We sought to identify the genetic variants associated with knee pain in 171,516 subjects from the UK Biobank cohort and replicate them using cohorts from 23andMe, the Osteoarthritis Initiative (OAI), and the Johnston County Osteoarthritis Study (JoCo).

**Methods:** We performed a genome-wide association study of knee pain in the UK Biobank, where knee pain was ascertained through self-report and defined as “knee pain in the last month interfering with usual activities”. A total of 22,204 cases and 149,312 controls were included in the discovery analysis. We tested our top and independent SNPs (*P* < 5 × 10^−8^) for replication in 23andMe, OAI, and JoCo, then performed a joint meta-analysis between discovery and replication cohorts using GWAMA. We calculated the narrow-sense heritability of knee pain using Genome-wide Complex Trait Analysis (GCTA).

**Results:** We identified 2 loci that reached genome-wide significance, rs143384 located in the *GDF5* (*P* = 1.32 × 10^−12^), a gene previously implicated in osteoarthritis, and rs2808772, located near *COL27A1* (*P* = 1.49 × 10^−8^). These findings were subsequently replicated in independent cohorts and increased in significance in the joint meta-analysis (rs143384: *P* = 4.64 × 10^−18^; rs2808772: *P* −11 = 2.56 × 10^−1^’). The narrow sense heritability of knee pain was 0.08.

**Conclusion:** In this first reported genome-wide association meta-analysis of knee pain, we identified and replicated two loci in or near *GDF5* and *COL27A1* that are associated with knee pain.

## Introduction

The knee supports body weight when walking, standing upright and bending. Knee pain describes a specific area of pain inside the knee or diffuse pain around knee area^1^. It is one of the most common musculoskeletal complaints that brings people to medical attention^2^. The knee pain experience varies from person to person and can present as a dull ache to a sharp, stabbing pain and from intermittent weight bearing pain to persistent pain^3^.

Knee pain is highly prevalent in older individuals, with ~50% of individuals over the age of 50 reporting an experience of knee pain within the past 12 months^4^. In one US general population cohort, knee pain prevalence has increased from 15.7% to 32.9% in females and from 8.7% to 27.7% in males between 1983 to 2005, regardless of knee osteoarthritis status^5^. In another estimate, the prevalence of knee osteoarthritis in the US increased from 8% in 1950s to 16% currently^6^. There are currently over 8 million patients suffering from knee osteoarthritis in the UK^7^. According to the Global Burden of Diseases 2016, osteoarthritis including knee osteoarthritis is the 12^th^ leading cause of years of life lived with disability (YLDs) globally^8^. It is estimated that ~ 50% of all people with knee osteoarthritis have reported knee pain symptoms and of those without knee osteoarthritis, 20% have reported knee pain^5^. There are many underlying mechanisms that can cause knee pain, including injuries, gout, and infection, as well as arthritis. Among these, osteoarthritis is the most common cause, particularly in people over the age of 50^5^. People with knee pain will experience progressive loss of knee function and declining quality of daily life, and display increasing dependence in daily activities^9^. Further, knee pain caused by osteoarthritis frequently accompanies pain in other joints such as hips and hands, which further reduces quality of life^10^. The disease has generated huge economic burdens to the health care systems across the world.

For example, although no figures exist specifically on knee osteoarthritis, the total direct cost of osteoarthritis as a whole in the UK in 2010 was around £1 billion and the corresponding total indirect cost of osteoarthritis in 2010 was over £3.2 billion^11^.

Epidemiological studies have suggested multiple risk factors for knee pain, including female sex, age, obesity, previous knee injuries, knee-straining work, and smoking^12^. Similar risk factors are reported in studies of knee osteoarthritis specifically, which also included kneeling and squatting as further risk factors^12,13^. With aging populations and increasing rates of obesity, the prevalence of knee pain is likely to increase. Psychological factors are also important risk factors of knee pain^14^. These environmental and lifestyle factors are likely to interact with genetic factors, and are important to understand in genetic association studies.

Genetic studies to date have focused on knee osteoarthritis, but not knee pain more generally. Studies in siblings have reported heritabilities for knee osteoarthritis as high as 0.62^15^. The genetic architecture of knee osteoarthritis was considered to follow an additive genetic model, involving multiple genes or loci but each with small effect size^16^. Candidate genes including *GDF5, COL9A1, IL1B, IL1RN, LRCH1, CLIP, TNA,* and *BMP2* have been reported to be associated with knee osteoarthritis^17–21^. Genome-wide association studies (GWAS) have also reported that the *GDF5, DVWA, HLA-DQB1, BTNL2, COG5, MCF2L, TP63, FTO, SUPT3H/RUNX2, GLN3/GLT8D1*, and *LSP1P3* genes contribute specifically to knee osteoarthritis^22–28^. Recently, Zengini et al reported 9 novel genetic loci associated with osteoarthritis based on 5 different osteoarthritis definitions according to the self-reported status questionnaire and the Hospital Episode Statistics data from the UK Biobank cohort^29^. However, these analyses did not specifically focus on the knee area.

To identify the genetic variants associated with knee pain, we conducted a GWAS using the large UK Biobank cohort. We defined knee pain as ‘knee pain in the last month interfering with usual activity’, based on the information available from the study questionnaire. Since there are no well-defined knee pain cohorts available, we chose to replicate our genome-wide significant findings on knee pain using three independent cohorts that defined osteoarthritis using either questionnaire data (i.e., 23andMe) or radiographic criteria (i.e., the Osteoarthritis Initiative (OAI) and the Johnston County Osteoarthritis Study (JoCo)).

As far as we know, this is the first GWAS on knee pain to screen for genetic variants. Similar approaches have been taken for headache and back pain using the UK Biobank cohort^30,31^.

## Method

### Participants and genetic information of the UK Biobank, 23andMe, OAI, and JoCo participants

Discovery cohort - UK Biobank: Over 500,000 people aged between 40 and 69 years were recruited by the UK Biobank cohort in 2006-2010 across England, Scotland and Wales. All participants provided informed consent that their health records could be accessed for research purposes. Further information about the UK Biobank cohort can be found at www.ukbiobank.ac.uk. Ethical approval was granted by the National Health Service National Research Ethics Service (reference 11/NW/0382).

DNA extraction and quality control (QC) were standardized and the detailed methods can be found at http://www.ukbiobank.ac.uk/wp-content/uploads/2014/04/DNA-Extraction-at-UK-Biobank-October-2014.pdf. The detailed QC steps can be found at http://biobank.ctsu.ox.ac.uk/crystal/refer.cgi?id=155580.

In July 2017, the UK Biobank released the genetic information (including directly genotyped genotypes and imputed genotypes) of 501,708 samples to all approved researchers. The detailed QC steps of imputation were described by Bycroft et al^32^.

Stage 1 replication cohort - 23andMe Inc: The 23andMe company is a privately held personal genomics and biotechnology company based in USA. It includes more than 1,500,000 genotyped subjects who have consented to participate in research. The DNA extraction from saliva and the QC of the genotyping and imputation results were all based on the company’s standardised procedures. Further methodological details can be found in the supplementary file of a previous publication^33^.

Stage 2 replication cohorts - 1. The OAI is a prospective longitudinal study designed to identify risk factors for the incidence and progression of symptomatic tibiofemoral knee osteoarthritis. Participants aged between 45 and 79 years were recruited at four different clinical sites in the USA. Details of the study protocol, including recruitment procedures and eligibility criteria are available on the OAI web site. (http://oai.epi-ucsf.org/datarelease/docs/StudyDesignProtocol.pdf). 2. The JoCo is an ongoing, community-based study of the occurrence of knee and hip osteoarthritis in African American and Caucasian residents, aged 45 years and above from Johnston County, North Carolina in the US. A total of 3,068 individuals were recruited at baseline. A detailed description of the cohort has been reported^34^.

Standard procedures of imputation and QC were applied when genotyping OAI and JoCo samples. The detailed description of the cohorts has been previously published.^28^

### Phenotypic information on knee pain of the UK Biobank, 23andMe, OAI, and JoCo

Discovery cohort - UK Biobank: We used a bespoke pain-related questionnaire adapted by the UK Biobank, which included the question: ‘in the last month have you experienced any of the following that interfered with your usual activities?’. The options were: 1. Headache; 2. Facial pain; 3. Neck or shoulder pain; 4. Back pain; 5. Stomach or abdominal pain; 6. Hip pain; 7. Knee pain; 8. Pain all over the body; 9. None of the above; 10. Prefer not to say. More than one option could be selected. (UK Biobank Questionnaire field ID: 6159)

The knee pain cases in this study were those who selected the ‘knee pain’ option for the above question, regardless of whether they had selected other options.

The controls in this study were those who selected the ‘None of the above’ option.

Stage 1 replication cohort - 23andMe, Inc: The 23andMe cohort used an online survey to determine the phenotypic status of osteoarthritis of all participants.

Cases were defined as those self-reported having been diagnosed or treated for osteoarthritis.

Controls were defined as those self-reporting as having not been diagnosed or treated for osteoarthritis.

The cohort included 253,880 cases and 1,286,245 controls.

Stage 2 replication cohorts - OAI and JoCo:

Cases: Knee osteoarthritis was evaluated with fixed-flexion posteroanterior radiographs for OAI samples and for JoCo participants, weight-bearing anteroposterior extended radiographs were taken during initial recruitments and fixed-flexion posteroanterior radiographs were taken during follow up. Cases were those with definitive knee osteoarthritis, defined as radiographic evidence of the presence of definite osteophytes and possible joint space narrowing (Kellgren-Lawrence grade ≥ 2) or total joint replacement in one or both knees. Controls were those having no or doubtful evidence of OA (KL grade = 0 or 1) in both knees at all available time points.

These definitions were previously used by Yau et al^28^. In the current study, there were 2,672 cases (2,014 from OAI and 658 from JoCo) and 1,776 controls (953 from OAI and 823 from JoCo).

### Statistical analysis

GWAS analysis: In the discovery stage, the BGENIE (https://jmarchini.org/bgenie/) was used as the main GWAS software recommended by the UK Biobank. Routine QC steps included removal of single nucleotide polymorphism (SNPs) with INFO scores less than 0.1, SNPs with minor allele frequency less than 0.5%, or SNPs that failed Hardy-Weinberg tests (*P* < 10^−6^). SNPs on the X and Y chromosomes and mitochondrial SNPs were also removed. We further removed data from individuals whose ancestry was not white British based on principal component analysis, those who were related to at least one other participant in the cohort (a cut-off value of 0.025 in the generation of the genetic relationship matrix), and those who failed QC. Association tests based on standard Frequentist association were performed using BGENIE adjusting for age, sex, body mass index (BMI), 9 population principal components, genotyping arrays, and assessment centres. A Chi-square test was used to test for gender differences between cases and controls. Age and BMI were compared using independent t testing in IBM SPSS 22 (IBM Corporation, New York). A *P* value less than 5 × 10^−8^ was considered to indicate a significant association. Independent SNPs were defined as those that were not correlated (r^2^<0.6) with any other significantly associated SNP. GCTA (https://cnsgenomics.com/software/gcta/#Overview) was used to calculate the narrow-sense heritability using a genomic relationship matrix calculated from genotyped autosomal SNPs.

In the replication stage, details of the identified significant and independent SNPs associated with knee pain in the discovery stage were sent to 23andMe Inc and the combined OAI and JoCo cohorts. The significant and independent SNPs were defined as those with *P* value < 5 × 10^−8^ and with linkage disequilibrium value r^2^ < 0.6. The 23andMe and the combined OAI and JoCo cohorts then extracted the summary statistics of these SNPs from their GWAS results, correspondingly.

23andMe performed GWAS on self-reported osteoarthritis in any joint using the logistic regression method assuming an additive genetic model for allelic effects adjusting for age, sex, 5 principal components, and 4 DNA chip platforms. Participants were restricted to a set of individuals who had >97% European ancestry, as determined through an analysis of local ancestry. A maximal set of unrelated individuals was chosen for the GWAS analysis using a segmental identity-by-descent (IBD) estimation algorithm. Further details can be found in the supplementary file of a previous publication^33^.

OAI and JoCo performed GWAS on radiographic knee osteoarthritis using logistic regression assuming an additive genetic model for allelic effects adjusting for age, sex, study site, and principal components. Summary statistics from both cohorts were then combined in a meta-analysis. Only participants of Caucasian origin were included in the GWAS study. Standard procedures were used to remove data from non-Caucasian individuals and related individuals. Further details can be found^28^.

The Genome-Wide Association Meta-Analysis (GWAMA) software (https://www.geenivaramu.ee/en/tools/gwama) was used to perform a meta-analysis combining the results from the UK Biobank, 23andMe, OAI, and JoCo cohorts.

GWAS-associated analysis: The FUMA web application was used as the main annotation tool, and a Manhattan plot and a Q-Q plot were also generated by this^35^. LocusZoom (http://locuszoom.org/) was used to provide regional visualization.

FUMA mainly provides 3 types of analysis: the gene analysis, the gene-set analysis and the tissue expression analysis. In gene analysis, summary statistics of SNPs were aggregated to the level of whole genes to test the associations between genes with the phenotype. In gene-set analysis, groups of genes sharing certain biological, functional or other characteristics were tested together to provide insight into the involvement of specific biological pathways or cellular functions in the genetic aetiology of a phenotype. The tissue expression analysis was based on GTEx (https://www.gtexportal.org/home/), which is integrated into FUMA.

To identify genetic correlations between knee pain and all other 234 complex traits, we used linkage disequilibrium score regression through LD Hub v1.9.0 (available at http://ldsc.broadinstitute.org/ldhub/) ^36^ The LD Hub estimates the bivariate genetic correlations of a phenotype with 234 traits using individual SNP allele effect sizes and the average linkage disequilibrium in a region. Those with *P* values less than 2.1 × 10^−4^ (0.05/234) were considered significant surviving Bonferroni correction for multiple testing.

## Results

### GWAS analysis results (Discovery cohort – UK Biobank)

A total of 501,708 UK Biobank participants were invited to respond to the pain questionnaire during the initial assessment visit (2006-2010). Among those who responded, 29,995 participants selected the ‘Knee pain’ option (cases), and 197,149 participants selected the ‘None of the above’ option (controls). After removing samples from non-British participants, those who were related with another individual in the cohort and those who failed QC, we identified 22,204 cases (12,062 males and 10,142 females) and 149,312 controls (71,480 males and 77,832 females) for the GWAS association analysis and there were 15,377,520 SNPs available for the GWAS analysis. The genomic control value (lambda) was 1.06.

Table I summarises the clinical characteristics of these cases and controls. There were statistical differences (P<0.001) in sex, age and BMI between cases and controls in the UK Biobank samples.

**Table I.** Clinical characteristics of knee pain and controls in the UK Biobank.

| Covariates | UK Biobank |  | *P* |
| --- | --- | --- | --- |
|  | Cases | Controls |  |
| Sex<br>(male:female) | 12062 :<br>10142 | 71,480 : 77,832 | <0.001 |
| Age (years) | 58.3 (7.64) | 56.9 (7.97) | <0.001 |
| BMI (kg/m<sup>2</sup>) | 28.6 (4.88) | 26.7 (5.00) | <0.001 |

BMI: body mass index A chi-square test was used to test the difference of gender frequency between cases and controls and an independent t test was used for other covariates. Continuous covariates were presented as mean (standard deviation).

We identified 2 SNP clusters that were associated with knee pain, with genome-wide significance (P < 5 × 10^−8^, Fig. 1, Table II). Four independent significantly associated SNPs within 2 clusters are shown in the Table II. All significantly associated SNPs (N=107) in the discovery stage are shown in Supplementary Table I.

**Table II.** Summary of the 4 independent and significant SNPs associated with knee pain in the *GDF5* and *COL27A1* regions.

| Chromosome | Gene | SNPID | Effective allele in all cohorts | Effective allele frequency Discovery cohort | *P* value (Beta) UK Biobank Discovery cohort | *P* value (Beta) 23andMe Replication stage 1 | *P* value (Beta) OAI-JoCo Replication stage 2 | *P* value meta-analysis | Beta meta-analysis | SE meta-analysis |
| --- | --- | --- | --- | --- | --- | --- | --- | --- | --- | --- |
| 9 | *LOC105376225 (near COL27A1)* | rs919642 | A | 73.2% | 2.29x10<sup>-8</sup> (0.007) | 4.84x10<sup>-12</sup> (0.028) | 0.88 (-0.008) | 1.20x10<sup>-13</sup> | 0.0093 | 0.0012 |
| 9 | *LOC105376225(near COL27A1)* | rs2808772 | A | 52.5% | 1.49x10<sup>-8</sup> (0.006) | 4.43x10<sup>-5</sup> (0.014) | 0.36 (0.041) | 2.56x10<sup>-11</sup> | 0.0073 | 0.0011 |
| 20 | *GDF5* | rs143384 | A | 60.0% | 1.32x10<sup>-12</sup> (-0.008) | 2.44x10<sup>-9</sup> (-0.021) | 0.01 (-0.12) | 4.64x10<sup>-18</sup> | -0.0095 | 0.0011 |
| 20 | *GDF5* | rs6120946 | A | 78.2% | 6.81x10<sup>-9</sup> (-0.008) | 0.00071 (-0.014) | 0.0074 (-0.16) | 3.52x10<sup>-11</sup> | -0.0088 | 0.0013 |

**Fig. 1.**
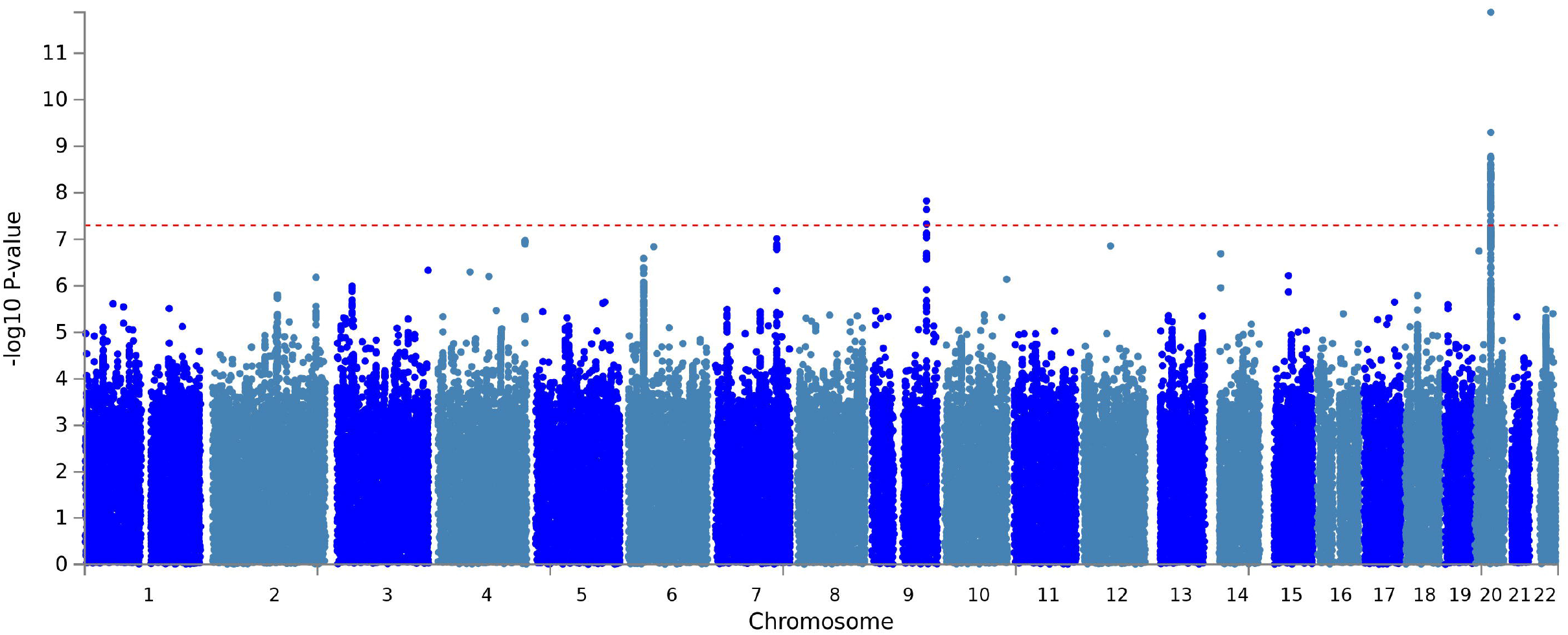
The Manhattan plot of the GWAS on knee pain using the UK Biobank cohort.

The most significantly associated SNP cluster was in the *GDF5* gene in the chromosome 20q11.22 region with a *P* value of 1.32 × 10^−12^ for rs143384 (A allele, odds ratio (OR): 0.992). The second most significantly associated cluster was in the *LOC105376225* gene (near the *COL27A1* gene) in chromosome 9 with a lowest *P* value of 1.49 × 10^−8^ for rs2808772 (A allele, OR: 1.006). The regional plots for loci in *GDF5* and *COL27A1* are shown in the Supplementary Fig. 1 and 2.

The Q-Q plot of the GWAS in the discovery stage is shown in the Supplementary Fig. 3. The SNP-based heritability of knee pain was 0.08 (standard error = 0.03).

### Replication stage results (23andMe, OAI, JoCo)

In the stage 1 replication, the *P* value of rs143384 was 2.44 × 10^−9^ in the 23andMe cohort and in the stage 2 replication, the *P* value of rs143384 was 0.01 in the combined OAI and JoCo cohorts. The *P* values of rs2808772 were 4.43 × 10^−5^ in the 23andMe and 0. 36 in the combined OAI and JoCo cohorts. (Table II)

### Meta-analysis results (UK Biobank, 23andMe, OAI, JoCo)

In the joint meta-analysis between the discovery and replication cohorts, the meta-analysis *P* values of all 4 independent and significant SNPs from 2 loci remained genome-wide significant and increased in significance. The meta-analysis results showed that the combined *P* values for rs143384 and rs2808772 were 4.64 × 10^−18^ (A allele, OR: 0.9905) and 2.56 × 10^−11^ (A allele, OR:1.007), respectively (Table II).

### Gene analysis, gene-set analysis and tissue expression analysis by FUMA

In the gene analysis, all the SNPs that are located within genes were mapped to 19,436 protein coding genes. *GDF5* demonstrated the most significant association, with a *P* value of 1.09 × 10^−11^. The 11 associated genes with *P* values less than 3 × 10^−6^ (0.05/19436) were *GDF5, UQCC1, CEP250, PODXL, C20orf173, SPAG4, MTMR3, ERGIC3, FBLN2, CPNE1, CDC42SE2.* The results are included in the Supplementary Table II.

In the gene-set analysis, a total of 10,894 gene sets were tested. The regulation pathway of breast_cancer_20q11_amplicon demonstrated a *P* value of 2.59 × 10^−8^ and this was the only gene set with a statistically significant association (*P* < 5 × 10^−6^ (0.05/10,894)). The top 10 gene sets from this analysis are shown in the Supplementary Table III.

In the tissue expression analysis, none of the tissue types demonstrated statistically significant associations (P < 0.001), either in the expression analysis of 30 general tissue types from multiple organs or in the 53 specific tissue types within some of these organs. See Supplementary Fig. 4 and 5.

### Genetic correlation analysis by LD Hub

We identified multiple significant and negative genetic correlations for knee pain with all other traits (Supplementary Table IV). The genetic correlations (r_g_) surviving multiple testing correction were: Years of schooling 2016 (r_g_ = −0.29, *P* = 4.97 × 10^−8^), College completion (r_g_ = −0.36, *P* = 6.55 × 10^−6^), Age of having first baby (r_g_ = −0.30, *P* = 1.92 × 10^−5^).

## Discussion

In the first reported GWAS of knee pain using the UK Biobank resource, we identified genome-wide significant variants in or near *GDF5* and *COL27A1,* which were subsequently replicated in osteoarthritis cohorts from the 23andMe, OAI, and JoCo cohorts. In addition, we found that knee pain was genetically and negatively correlated with a number of socioeconomic factors such as years of schooling and college completion.

The generic pain question used by the UK Biobank is useful as a screening tool and a useful step to test whether heterogeneous pain phenotypes (such as knee pain) have genetic components at all. We have successfully used the same question to identify the genetic variants of broadly-defined headache, and our findings were similar to those for well-defined migraine phenotypes^30,39^. The benefit of using UK Biobank on heterogeneous phenotypes will allow researchers to overcome potential issues with reduced power due to heterogeneity by using very large numbers to cut through the statistical noise.

In this GWAS, we have identified 2 loci for knee pain. The top locus was in the *GDF5* gene in chromosome20q11.2 with a lowest *P* value of 1.32 × 10^−12^ for rs143384 while the locus itself was 140kb long spanning from the *UQCC1* gene to the *GDF5* gene containing 104 significant SNPs (Supplementary Table I). The *GDF5* gene encodes a secreted ligand of the transforming growth factor-beta (TGF-beta) superfamily of proteins^40^. This protein not only regulates the development of numerous tissue and cell types, but also promotes the maintenance and repair of synovial joint tissues, particularly cartilage and bones^40,41^. Mutations in the gene can cause cartilage or bone related disorders such as chondrodysplasia, acromesomelic dysplasia, and brachydactyly, suggesting a protective role in skeletal development^40^. The *GDF5* gene has been repeatedly reported to be associated with osteoarthritis through genetic studies^17,27^. Functional studies have suggested that knee morphology is profoundly affected by Gdf5 absence in mice models, and downstream regulatory sequences mediate its effects by controlling Gdf5 expression in knee tissues^42^. It was also suggested that osteoarthritis susceptibility mediated by variants in the *GDF5* gene was not restricted to cartilage but joint wide^43^. Recently, Terence et al combined transgenetic mice model with population genetic analyses in humans to identify a GDF5 enhancer that influences human growth and osteoarthritis risk^44^. Overall, there is sufficient and solid biological evidence relating the *GDF5* gene with knee osteoarthritis, and we assume that this finding is due to detection of knee pain caused by osteoarthritis, rather than other pathologies.

The second SNP cluster was in the *LOC105376225* area (which is next to the *COL27A1* gene) in chromosome 6 with a lowest *P* value of 1.49 × 10^−8^ for rs2808772. There have been no specific studies published about *LOC105376225* and its relationship with knee pain or knee osteoarthritis. However, the neighbouring *COL27A1* gene is clearly a good candidate gene. This gene encodes a member of the fibrillar collagen family, and plays a role during the calcification of cartilage and the transition of cartilage to bone^45^. Mutations in the *COL27A1* gene have been reported to be associated with the Steel syndrome. This syndrome is characterized by bone changes such as bilateral hip and radial head dislocations, short stature, characteristic facies, fusion of carpal bones, scoliosis, *pes cavus,* and cervical spine anomalies^46^. Further, the gene was reported to be associated with knee osteoarthritis in the first stage, but did not replicate in the second stage in a recent GWAS study on knee osteoarthritis^28^. Thus, our large study on knee pain has suggested that the *COL27A1* gene might play a role in the knee area. Importantly, polymorphisms in the gene have been associated with tendinopathy around the ankle joint^47^. The concordance between radiographically defined knee osteoarthritis and knee pain is quite poor, with between 15% and 81% of patients diagnosed by radiographic methods having pain symptoms^48^. It is therefore likely that many people reporting knee pain have pain that is not bone or cartilage related, but tendon related.

Our study focused on knee pain as a broad phenotype and the genes that we identified and replicated are suggested to be related to knee osteoarthritis. This suggests that the phenotype we chose was genetically similar to the phenotype of knee osteoarthritis. The relationship between knee pain and knee osteoarthritis deserves further investigation. Studies have shown that people with end-stage knee osteoarthritis all presented with knee pain^49^, but this might not be the case for early stage knee osteoarthritis. As described above, 20% of knee pain was not caused by knee osteoarthritis and only 50% of knee osteoarthritis patients with radiographic evidence had knee pain symptoms^5^.

The gene-analysis by FUMA also supports our finding that *GDF5* was the strongest gene for knee pain. The gene-set analysis by FUMA revealed that the regulation pathway of Nikolsky_breast_cancer_20q11_amplicon signaling was associated with the phenotype we use. We noticed that *GDF5* and this amplicon were both located in chromosome 20q11 area and it was reported that GDF5 protein regulates TGF-beta dependent angiogenesis in breast carcinoma MCF-7 Cells^50^.

The SNP-based heritability for knee pain was 0.08 in our study, which is the first report of its kind. We identified that knee pain was genetically and negatively correlated with a number of phenotypes such as years of schooling, college completion and age of having first baby. This means that those with more years of schooling, those with completed college education, and those who were older when they had their first baby were less likely to report current troublesome knee pain. These factors could be related to lifestyle and occupation.

Using the CaTS power calculator (http://csg.sph.umich.edu/abecasis/cats/), we had 80% power to identify SNP associations with a significance level of 5 × 10^−8^, based on 22,204 cases and 149,312 controls, assuming an additive model, a minor disease allele frequency of 0.20, a genotypic relative risk of 1.06, and an estimated prevalence of knee pain in the general population of 0.2.

In this study, we used different but similar phenotypes in discovery and replication stages. We defined cases and controls based on the responses by UK Biobank participants to a specific pain question. This question focused on knee pain occurrence, sufficient to cause interference with activities, during the previous month. The question does not ask information of the severity and frequency and the exact area of knee pain. Therefore, our phenotyping should be considered as widely defined. The situation was similar for the 23andMe cohort, in which disease status was also self-reported via survey and self-reported osteoarthritis in the 23andMe cohort was not specific to the knee. Self-reported knee pain has been widely used in other studies as well, though not for studies of genetic associations^37,38^. The OAI and JoCo assessed radiographic evidence of knee osteoarthritis but with limited sample size.

In conclusion, we have identified 2 loci (*GDF5* and *COL27A1*) for knee pain in a GWAS using the UK Biobank resource and replicated them in the 23andMe, OAI, and JoCo cohorts. In addition, we found several significant and negative genetic correlations between knee pain and a number of educational phenotypes, suggesting that the genetic aetiology of knee pain may also be related to these traits.

## Acknowledgements

We would like to thank all participants of the UK Biobank, 23andMe and the OAI and JoCo cohorts who have provided necessary genetic and phenotypic information. The current study was conducted under approved UK Biobank data application number 4844.

Members of the 23andMe Research Team: Michelle Agee, Babak Alipanahi, Robert K. Bell, Katarzyna Bryc, Sarah L. Elson, Pierre Fontanillas, Nicholas A. Furlotte, Barry Hicks, David A. Hinds, Karen E. Huber, Ethan M. Jewett, Yunxuan Jiang, Aaron Kleinman, Keng-Han Lin, Nadia K. Litterman, Jennifer C. McCreight, Matthew H. McIntyre, Kimberly F. McManus, Joanna L. Mountain, Elizabeth S. Noblin, Carrie A.M. Northover, Steven J. Pitts, G. David Poznik, J. Fah Sathirapongsasuti, Janie F. Shelton, Suyash Shringarpure, Chao Tian, Joyce Y. Tung, Vladimir Vacic, Xin Wang, Catherine H. Wilson.

## Author contributions

WM organised project, drafted the paper and contributed to the analysis. MA performed the main UK Biobank GWAS analysis. CP, BM, MY provided essential comments. KR, JJ, BM, RJ and MY provided the OAI and JoCo GWAS summary statistics on knee osteoarthritis. JS and AA performed replication in the 23andMe cohort. AM and BS organised the project and provided comments.

## Data availability

The summary statistics of the UK Biobank results on knee pain can be shared upon request to non - commercial researchers.

