## Supplementary material for "A genome-wide association study identifies that the GDF5 and COL27A1 genes are associated with knee pain in UK Biobank (N = 171, 516)": Sup Table 2

| Gene | Chromosome | Start | Stop | Number of SNPs | Z-STAT | *P* |
| --- | --- | --- | --- | --- | --- | --- |
| *GDF5* | 20 | 34021145 | 34042568 | 66 | 6.6939 | 1.09E-11 |
| *UQCC1* | 20 | 33890369 | 33999944 | 358 | 6.6671 | 1.30E-11 |
| *CEP250* | 20 | 34042985 | 34099804 | 189 | 6.3487 | 1.09E-10 |
| *PODXL* | 7 | 131185021 | 131242976 | 297 | 5.2024 | 9.84E-08 |
| *C20orf173* | 20 | 34111014 | 34117481 | 18 | 5.1655 | 1.20E-07 |
| *SPAG4* | 20 | 34203814 | 34208971 | 15 | 5.0917 | 1.77E-07 |
| *MTMR3* | 22 | 30279144 | 30426855 | 623 | 4.8555 | 6.00E-07 |
| *ERGIC3* | 20 | 34129770 | 34145405 | 50 | 4.8223 | 7.10E-07 |
| *FBLN2* | 3 | 13573824 | 13679922 | 701 | 4.6333 | 1.80E-06 |
| *CPNE1* | 20 | 34213953 | 34252878 | 143 | 4.5625 | 2.53E-06 |
| *CDC42SE2* | 5 | 130581186 | 130734140 | 508 | 4.5372 | 2.85E-06 |

Supplementary Table 2 the 11 associated genes of the gene analysis by FUMA
