## Supplementary material for "A genome-wide association study identifies that the GDF5 and COL27A1 genes are associated with knee pain in UK Biobank (N = 171, 516)": Sup Table 3

| Full name of the pathways | Number of genes | Beta | Beta-STD | SE | *P* |
| --- | --- | --- | --- | --- | --- |
| Curated_gene_sets:nikolsky_breast_cancer_20q11_amplicon | 31 | 1.44 | 0.0605 | 0.268 | 3.76x10^-8^ |
| GO_cc:go_presynaptic_active_zone | 25 | 0.765 | 0.0288 | 0.187 | 2.16x10^-5^ |
| Curated_gene_sets:biocarta_tid_pathway | 13 | 0.776 | 0.0211 | 0.222 | 0.000237 |
| GO_cc:go_anchored_component_of_external_side_of_plasma_membrane | 19 | 0.697 | 0.0229 | 0.202 | 0.000278 |
| Curated_gene_sets:figueroa_aml_methylation_cluster_7_dn | 5 | 1.48 | 0.025 | 0.433 | 0.000317 |
| GO_cc:go_intrinsic_component_of_external_side_of_plasma_membrane | 23 | 0.615 | 0.0222 | 0.183 | 0.000402 |
| Curated_gene_sets:traynor_rett_syndrom_dn | 15 | 0.746 | 0.0218 | 0.229 | 0.000548 |
| Curated_gene_sets:kegg_tgf_beta_signaling_pathway | 75 | 0.32 | 0.0208 | 0.0986 | 0.000598 |
| GO_bp:go_embryonic_camera_type_eye_development | 32 | 0.537 | 0.0229 | 0.17 | 0.000797 |
| GO_bp:go_mating_behavior | 22 | 0.589 | 0.0208 | 0.188 | 0.000877 |

Supplementary Table 3: the top 10 gene sets generated by FUMA

STD: Standard deviation

SE: Standard error
