## Supplementary material for "A genome-wide association study identifies that the GDF5 and COL27A1 genes are associated with knee pain in UK Biobank (N = 171, 516)": Sup Fig 2

### Slide 1
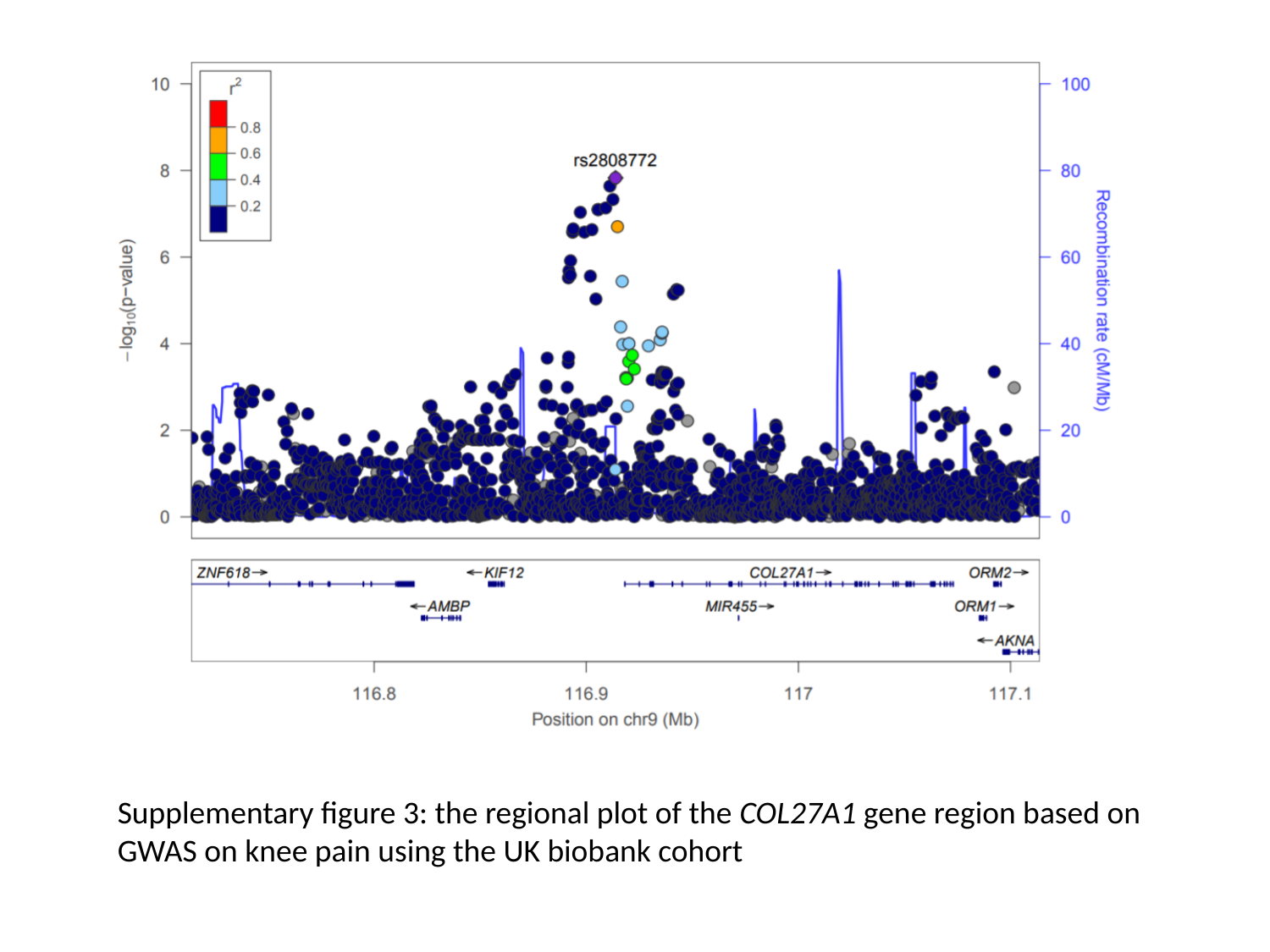

Supplementary figure 3: the regional plot of the COL27A1 gene region based on GWAS on knee pain using the UK biobank cohort
