## Supplementary figures and images for "A genome-wide association study identifies that the GDF5 and COL27A1 genes are associated with knee pain in UK Biobank (N = 171, 516)"

### Sup Fig 3

#### Slide 1
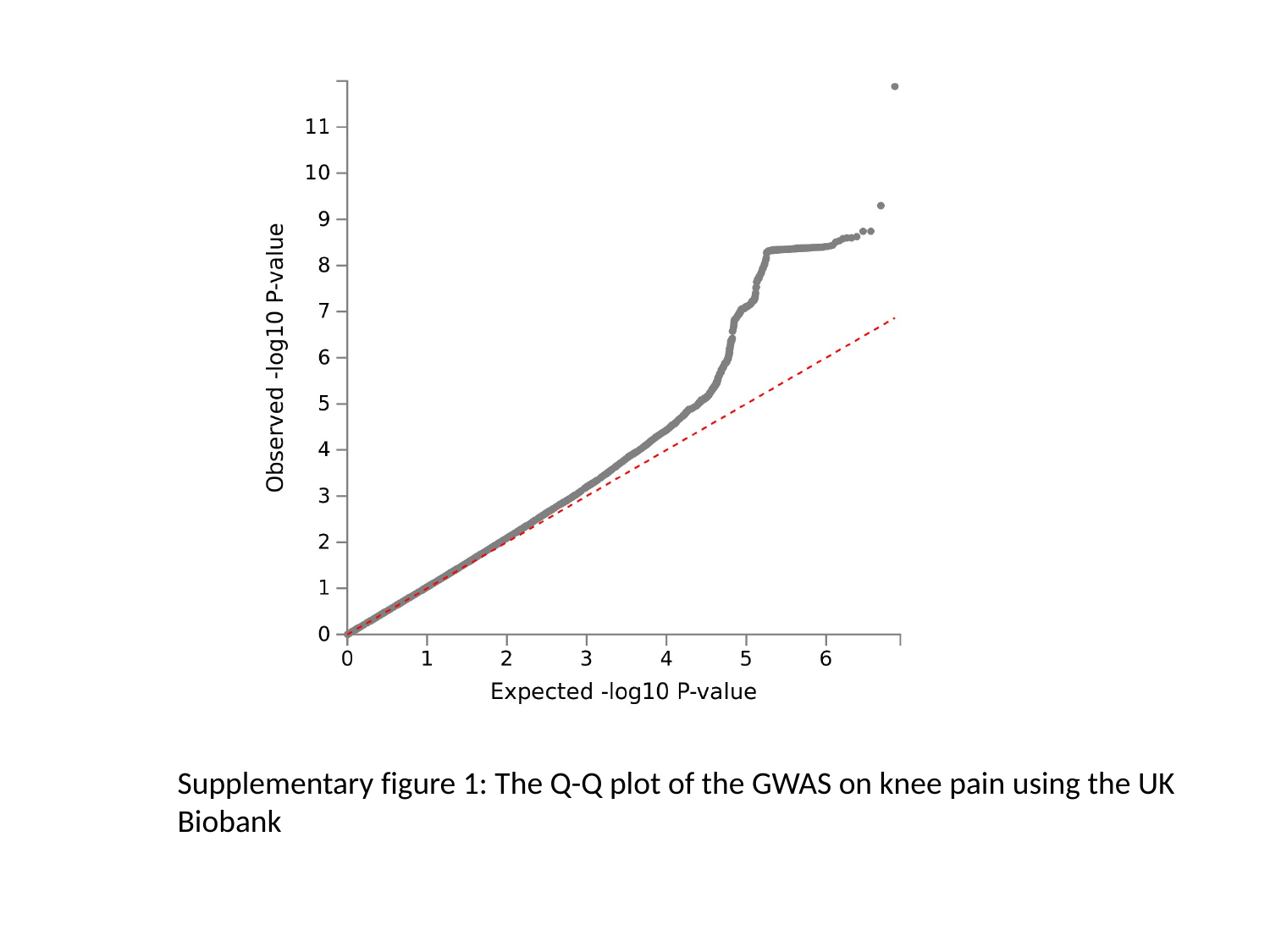

Supplementary figure 1: The Q-Q plot of the GWAS on knee pain using the UK Biobank

### Sup Fig 4

#### Slide 1
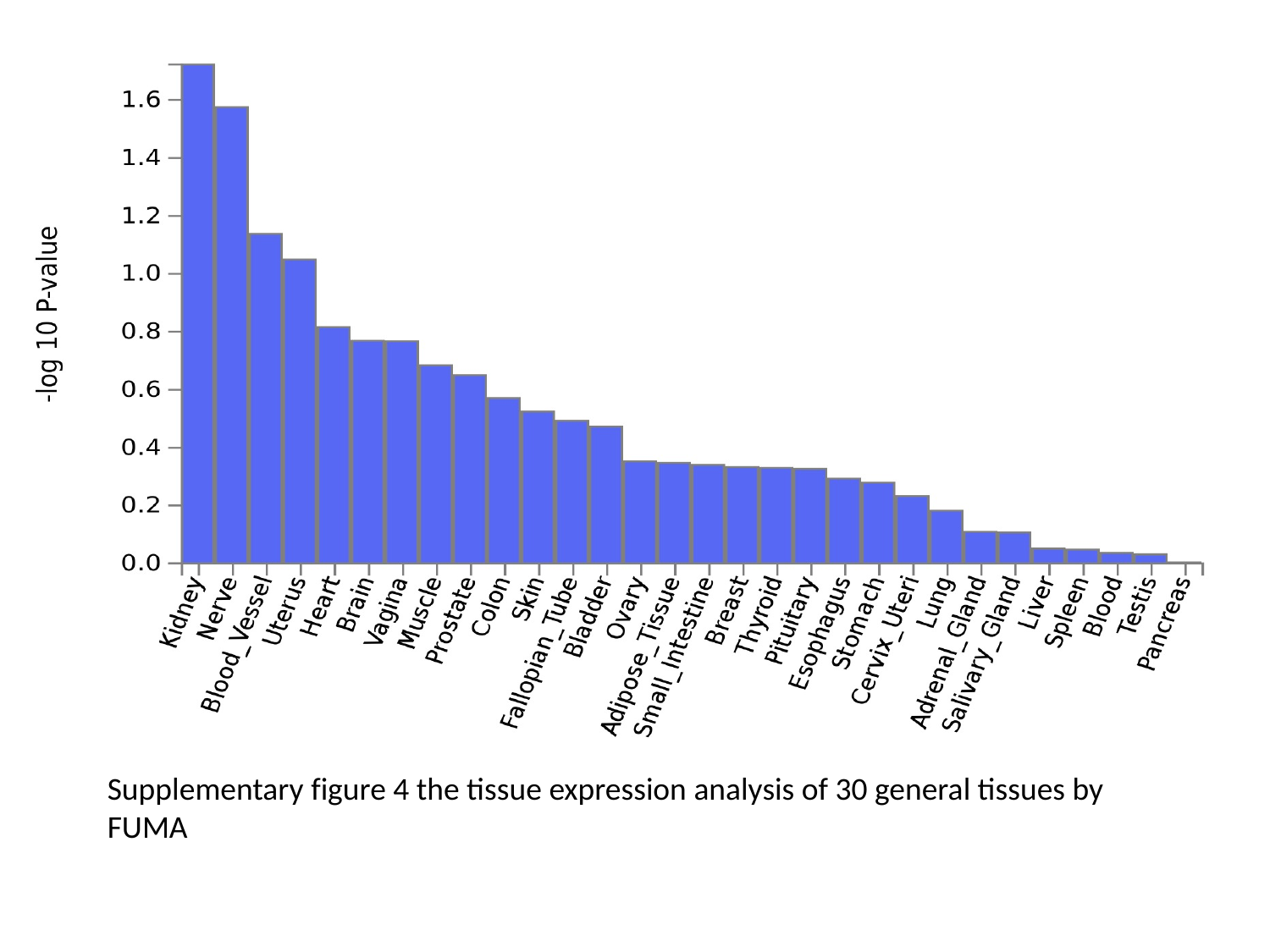

Supplementary figure 4 the tissue expression analysis of 30 general tissues by FUMA

### Sup Fig 5

#### Slide 1
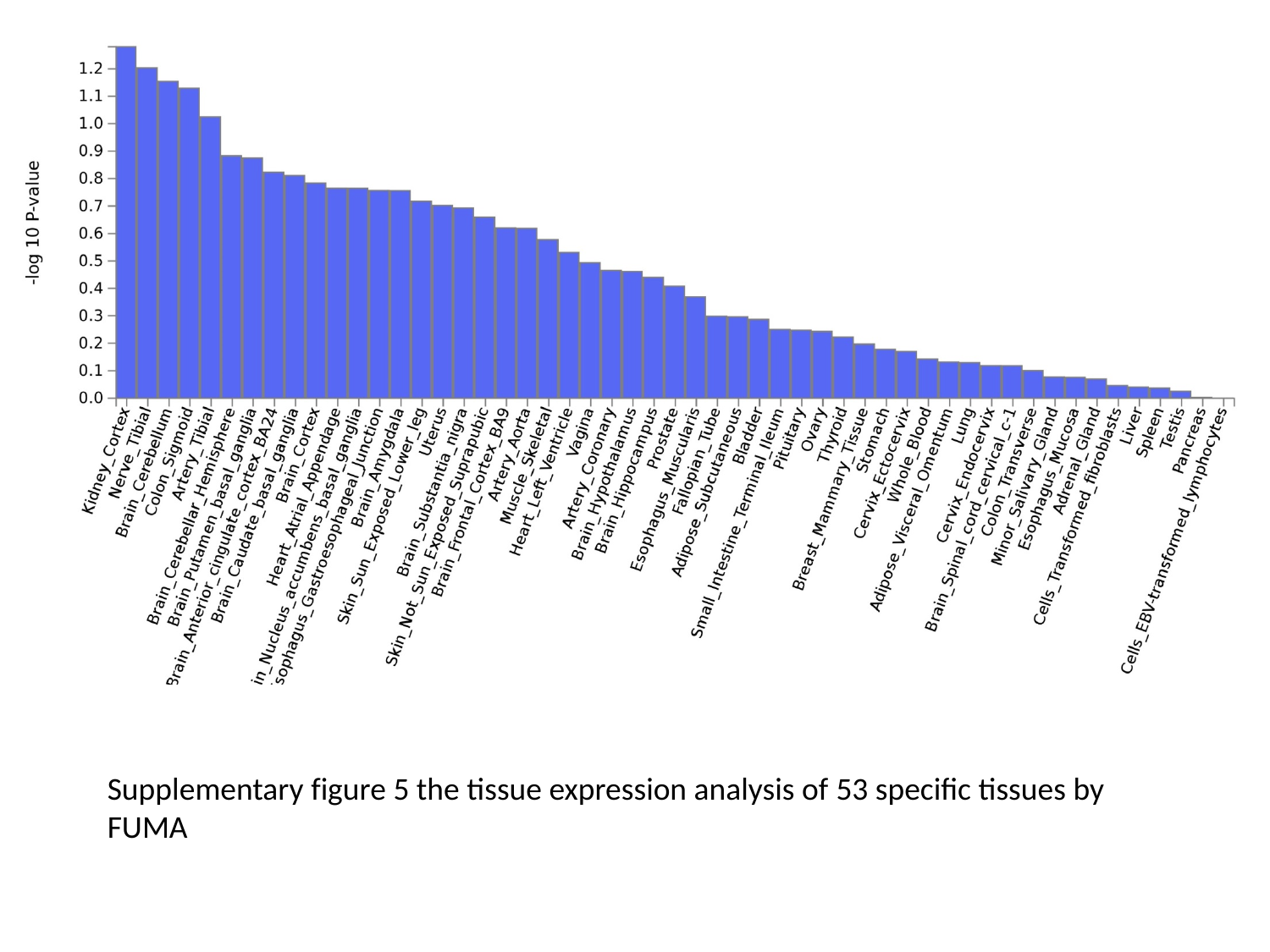

Supplementary figure 5 the tissue expression analysis of 53 specific tissues by FUMA
